# Transcranial Direct Current Stimulation over the Posterior Parietal Cortex Increases Non-target Retrieval during Visual Working Memory

**DOI:** 10.1101/2024.06.17.599451

**Authors:** Shengfeng Ye, Menglin Wu, Congyun Yao, Gui Xue, Ying Cai

## Abstract

Accurate visual working memory (VWM) requires precise content-context binding. Previous studies have revealed a close relationship between the posterior parietal cortex (PPC) and feature binding during VWM, this study further examined their causal relationship through three transcranial direct current stimulation (tDCS) experiments. In Experiment 1 (*N* = 57), participants underwent three sessions of tDCS separately, including PPC stimulation, occipital cortex stimulation, and sham stimulation, and completed a series of delayed estimation tasks for orientations before and after stimulation. Results showed that tDCS over PPC selectively prolonged recall response time (RT) and increased the probability of non-target responses (a.k.a. failure of feature binding). In Experiment 2 (*N* = 29), combining metacognition estimation during the task, we further investigated whether the effects of PPC stimulation on RT and increased probability of non-target responses were attributed to more mis-binding (i.e., participants self-reported "remembered" in non-target responses) or informed guessing (participants self-reported "forgotten" in non-target responses). We replicated the main findings in Experiment 1, and we also observed greater tDCS effects of PPC on RT in informed guessing trials than mis-binding trials while comparable effects on non-target response rates in these two types of trials. In Experiment 3 (*N* = 28), we then examined whether the effects of tDCS over PPC specifically influenced the memory retrieval process by using a change detection task. We found that PPC stimulation did not influence the recognition RT or accuracy. Together, this study provides causal evidence supporting the involvement of PPC in feature binding during VWM retrieval.

**Significance Statement:** Visual working memory (VWM) enables humans to temporarily store and process visual information, which requires accurate binding of items to their unique context. Accumulating studies posited that the posterior parietal cortex (PPC) is closely related to this binding process, the current study further examined their causal relationship. Through three strictly within-subject well-designed non-invasive neural stimulation experiments, we found that PPC stimulation selectively increased response time (RT) and binding error during VWM. Moreover, we found these changes were modulated by individual metacognition and only occurred during memory recall instead of recognition. Together, our results provided strong evidence that PPC is causally involved in the binding process during visual working memory retrieval.

## Introduction

Visual working memory (VWM), a process of storing and processing visual information temporarily, is an essential basis of higher-level cognitive processes. Precise content-context binding is critical for multi-item VWM. The latest interference model indicates that VWM capacity is mainly limited by the binding interference between the content of multiple memory items and their specific context information (Oberauer & Lin, 2017). Moreover, impaired binding ability is an early symptom in a wide range of neurodegenerative disorders (Kirova et al., 2015; Mayer et al., 2012). Thus, understanding the mechanism underlying content-context binding in VWM is always a critical theoretical question.

Increasing studies have demonstrated the relationship between the posterior parietal cortex (PPC) activity and feature binding during VWM. In an early study, patients with right PPC lesions exhibited a decreased accuracy in tasks requiring multi-item bindings (Ashbridge et al., 1999) and usually reported illusory feature conjunctions during memory recall tests (Braet & Humphreys, 2009). Although earlier functional magnetic resonance imaging (fMRI) and electroencephalograph (EEG) studies have revealed that PPC activity was tightly correlated with the number of items kept during VWM (Todd & Marois, 2004), recent fMRI studies showed that, holding the memorized items constant, PPC activity was higher when content-context binding demands increased, suggesting that PPC played a closer role in the binding process (Cai et al., 2020; Gosseries et al., 2018).

Besides, modeling studies have advanced the relationship between PPC activity and feature binding to the individual level trial level. For example, Bays et al. (2009) proposed a multiple-component mixture model to estimate individuals’ probability of non-target responses, which reflected the failure of binding in a delayed estimation recall task. Accordingly, Cai et al. (2020) found that the neural decoding strength of the context information (i.e., location) in PPC could predict rates of non-target responses. Moreover, Schneegans and Bays (2016) proposed a novel model that could estimate the probability of non-target recalls even at the trial level. By combining this model and metacognitive reports, researchers further identified two types of non-target responses: high-confidence mis-binding and low-confidence informed guessing (van den Berg et al., 2017). Based on this view, Mallett et al. (2022) observed that participants responded with high confidence in about three-quarters of non-target trials in a delayed estimation task. More importantly, they found that non-target items could be decoded from PPC activity since early maintenance. Unfortunately, this study did not compare the differences between the two types of non-target responses, thus the involvement of PPC in these two types of non-target responses was still unclear.

Transcranial direct current stimulation (tDCS) is an important tool to explore the causal relationship between neural activity and cognitive processes. Currently, tDCS research testing the relationship between PPC and VWM has focused on memory capacity, while the results were quite controversial (Dumont et al., 2021; Jiang et al., 2023; Li et al., 2017; Wang et al., 2019). For these inconsistencies, meta-analyses have pointed out that different control groups could be an important factor (Dedoncker et al., 2016; Hill et al., 2016). Meanwhile, more specifically, researchers have also proposed some individual differences modulating tDCS effects, such as VWM baseline capacity (Hsu et al., 2014; Tseng et al., 2012), remember-subset or remember-all strategy (Wang et al., 2020), and biorhythms during stimulation (i.e., morning vs. afternoon; (Salehinejad et al., 2019; Salehinejad et al., 2023; Salehinejad et al., 2021). In sum, few tDCS studies have examined the causal relationship between PPC and feature binding, and to answer this question, it is critical to use control irrelevant variables and fully explore the potential individual differences that affected tDCS effects.

In this study, we systematically investigated the causal relationship between PPC and feature binding in VWM through three tDCS experiments. In Experiment 1, combined with the three-factor mixture model, we examined the effect of tDCS over PPC on binding process in a delayed estimation task and explored how individual differences in capacity and recall strategies affected the tDCS effect. In Experiment 2, we tried to replicate the results of Experiment 1 in an independent sample as well as further investigated the involvement of PPC in two types of non-target responses (i.e., mis-binding and informed guessing), by integrating remembered-forgotten self-reports during the task. In Experiment 3, we examined whether the tDCS effects over PPC on binding process specifically during memory retrieval by using a change detection task (e.g., recognition) instead of a delayed estimation task (e.g., recall). In this case, if tDCS changed VSTM maintenance, similar effects should be observed in both tasks; otherwise, no such effect would be detected in the recognition task.

## Experiment 1

### Participants

Fifty-eight university students participated in Experiment 1 (34 females; *M* = 20.10 years, *SD* = 1.40). All participants were right-handed, had normal or corrected-to-normal vision, and reported no history of neurological or psychiatric disorders. Before the experiment, participants signed a written informed consent required by the institutional review board of the Department of Psychology and Behavioral Sciences, [Author University]. Participants received monetary compensation for participation (¥30 per hour).

### Materials and methods

We modified a delayed estimation task for orientations from previous studies (Wang et al., 2019, 2020). In each trial, after a 500 ms white central fixation (0.75° × 0.15°), an array of 6 or 8 white, randomly oriented bars (2° × 0.3°) were presented for 200 ms. All bars were uniformly presented on an invisible circle (centered on the screen, radius of 6°), and orientations of these stimuli were randomly selected from 10° to 170° with at least 10° apart. After a 1 s delay, participants were instructed to recall the orientation of the bars at the probed position as precisely and fast as possible, by moving the mouse and clicking the left bottom on the white circular probe (radius of 2°). The maximum response time was 8 s. Participants completed the task before and after each tDCS session for approximately 20 minutes. Each session consisted of 240 trials which were divided into 4 blocks, with two memory loads randomly mixed. Participants completed 24 practice trials in each memory load condition before the formal experiment to become familiar with the task.

Experimental stimuli were displayed on a 17-inch color screen running MATLAB 2019a (MathWorks; US) and the Psychophysics Toolbox 3.0.12 (Brainard & 1997). Participants were seated 60 cm away from the monitor (resolution, 1920×1080; refresh rate, 60 Hz) in a quiet room and were instructed to keep their eyes fixed on the center of the screen throughout the experiment. At the end of the experiment, participants completed a 5-point scale to evaluate how often they use remember-all strategy or remember-subset strategy under different memory loads (1 = always “remember-all”; 5 = always “remember-subset”).

### tDCS setup

Participants underwent three tDCS sessions, including PPC stimulation, occipital cortex (OCC) stimulation, and sham stimulation. The order of stimulations was counterbalanced across participants, and three stimulations were performed separately with an exactly 48-hour interval to minimize potential carryover effects and influences of circadian rhythms (Figure 1B). tDCS was delivered by the DC-STIMULATOR MC device (NeuroConn; Germany) using a pair of plastic electrodes (5×7 cm^2^) in a saline-soaked synthetic sponge. In PPC stimulation, the anodal electrode was placed over P4 according to the International 10-20 electroencephalogram (EEG) electrode system (Hsu et al., 2014; Li et al., 2017; Wang et al., 2019, 2020), the reference electrode was placed over the left cheek. Then a 20-minute, 2 mA current was applied, with a linear fade in and fade out of 30 s which could minimize the uncomfortable feelings of sudden current changes. In OCC stimulation, the only difference is the anodal electrode was placed over Oz (Li et al., 2017; Makovski & Lavidor, 2014). For the sham condition, the anodal electrode was randomly placed over P4 or Oz, and the stimulation was only delivered within the first and last 30 seconds to simulate the itching feelings during active stimulation. During all stimulations, participants sat and took a rest. The current density distributions for tDCS settings were presented using COMETS, an open-source toolbox based on MATLAB (http://www.cometstool.com; Lee et al., 2017; Figure 1C).

**Figure 1.**
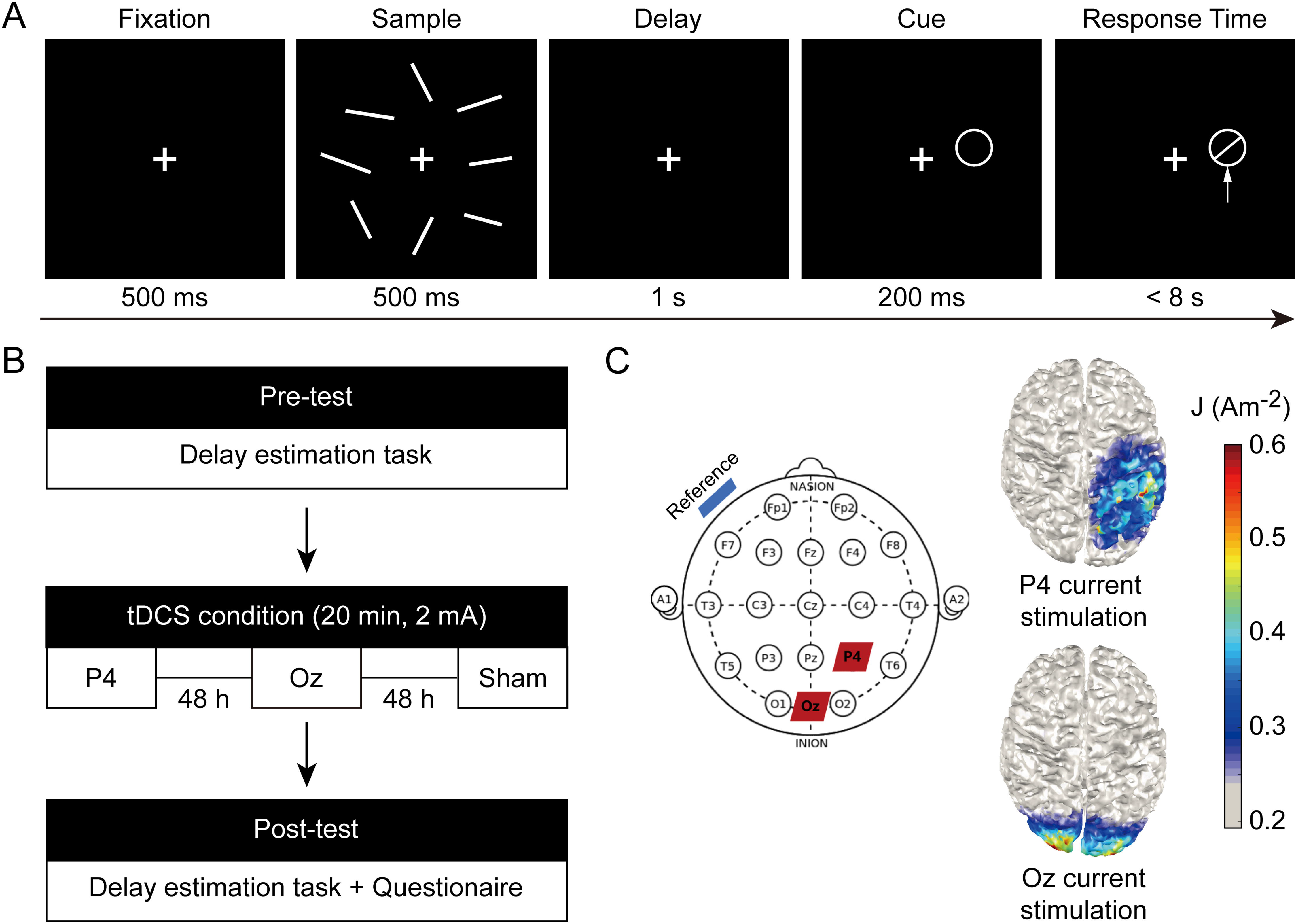
(A) Schematic diagrams of the delay estimation task. (B) tDCS procedures. (C) The placement of tDCS electrodes (left) with red patches representing the anodal electrode (P4 and Oz) and a blue patch representing the reference electrode (left cheek) and cortical current density distributions from an overhead view (right).

### Data analysis

#### Estimations of behavioral performance

We calculated the response time (RT) and the recall error under different memory loads. RT was defined as the duration between cue onset and response confirmation, and the recall error was obtained as the angular distance between the reported orientation and the targeted orientation. Then, we adopted the three-factor mixture model (Bays et al., 2009) and fit the distribution of raw errors to obtain the precision of target responses (κ), the probability of target responses (*p*T), the probability of non-target responses (*p*NT), and the probability of guess responses (*p*U). Among these measures, a higher *p*NT indicated a higher probability of content-context binding errors. Memory capacity was calculated as *p*T × memory load according to previous research (Zhang & Luck, 2008). One participant was excluded due to poor working memory performance (capacity < 1). Then, 57 participants were included for further analysis.

#### Estimations of tDCS effects

To examine tDCS effects, we first compared the behavioral parameters between active stimulation (PPC or OCC stimulation) and sham stimulation separately. For each behavioral parameter, we conducted a three-way repeated-measures analysis of variance (ANOVA) of stimulation condition (active vs. sham), test time (pre-test vs. post-test), and memory load (set size 6 vs. set size 8). If interaction effects were significant, we then performed two-way repeated-measures ANOVAs investigating the tDCS effects under each load. Furthermore, if any tDCS effect was significant in active stimulation, we tested whether this effect was region-specific by comparing tDCS effects over PPC and OCC with repeated-measures ANOVAs as well as calculating the Pearson correlation between PPC and OCC tDCS effects (i.e., the behavioral changes before and after stimulation). We reported *p* values, effect size, and Bayesian factor (BF) for all statistical analyses.

#### Examination of influence factors of tDCS effects

For significant tDCS effects at the group level, we examined the potential influences of individual working memory capacity and recall strategy scores on tDCS effects via Pearson correlation analyses. Because the capacity measures (i.e., *p*T × each memory load) were highly correlated in three pre-stimulation sessions and two memory loads (*r*s > 0.379, *p*s < 0.01, BF_10s_ > 10.263), we averaged them as the individual capacity performance. Similarly, the recall strategy scores were obtained in each memory load, which was not correlated (*r* = 0.613, *p* < 0.001, BF_10_ > 1000). Then, we averaged them as the individual memory strategy if the tDCS effects were significant in both set sizes, while only using the recall strategy score in the specific memory load (i.e., set size 8) if the tDCS effect was load-specific.

### Results

#### PPC stimulation prolonged recall response time

For tDCS effects over PPC in RT, the interaction effect of stimulation condition, test time, and memory load was not significant (*F*_(1,56)_ = 0.595, *p* = 0.444, *η_p_*^2^ = 0.011, BF_10_ = 0.205), whereas the interaction between stimulation condition and test time was significant (*F*_(1,56)_ = 6.228, *p* = 0.016, *η_p_*^2^ = 0.100, BF_10_ = 3.366). For both memory loads, we observed significant interactions between stimulation condition and test time (*F*s > 5.179, *p*s < 0.027, *η_p_*^2^s > 0.085, BF_10_s > 2.309). Simple effect tests revealed that, although both PPC stimulation and sham stimulation lead to significantly reduced RT (*t*s > 5.406, *p*s < 0.001, Cohen’s *d*s > 0.716, BFs_10_ > 1000), the decrease was smaller in tDCS over PPC compared with that in sham (i.e., a relatively longer RT after tDCS over PPC, Figure 2A. left). Meanwhile, at the individual level, the tDCS effects on prolonged RTs between two memory loads were highly correlated (*r*_(55)_ = 0.864, *p* < 0.001, BF_10_ > 1000; Figure 2B, top). Thus, we averaged RT changes in two memory loads to index the tDCS effects on RT in the following analysis. For tDCS effect over OCC in RT, on the contrary, our results revealed no three-way nor two-way interactions between test time and stimulation conditions (*F*s < 2.622, *p*s > 0.111, *η_p_*^2^s < 0.045, BF_10_s < 0.458).

**Figure 2.**
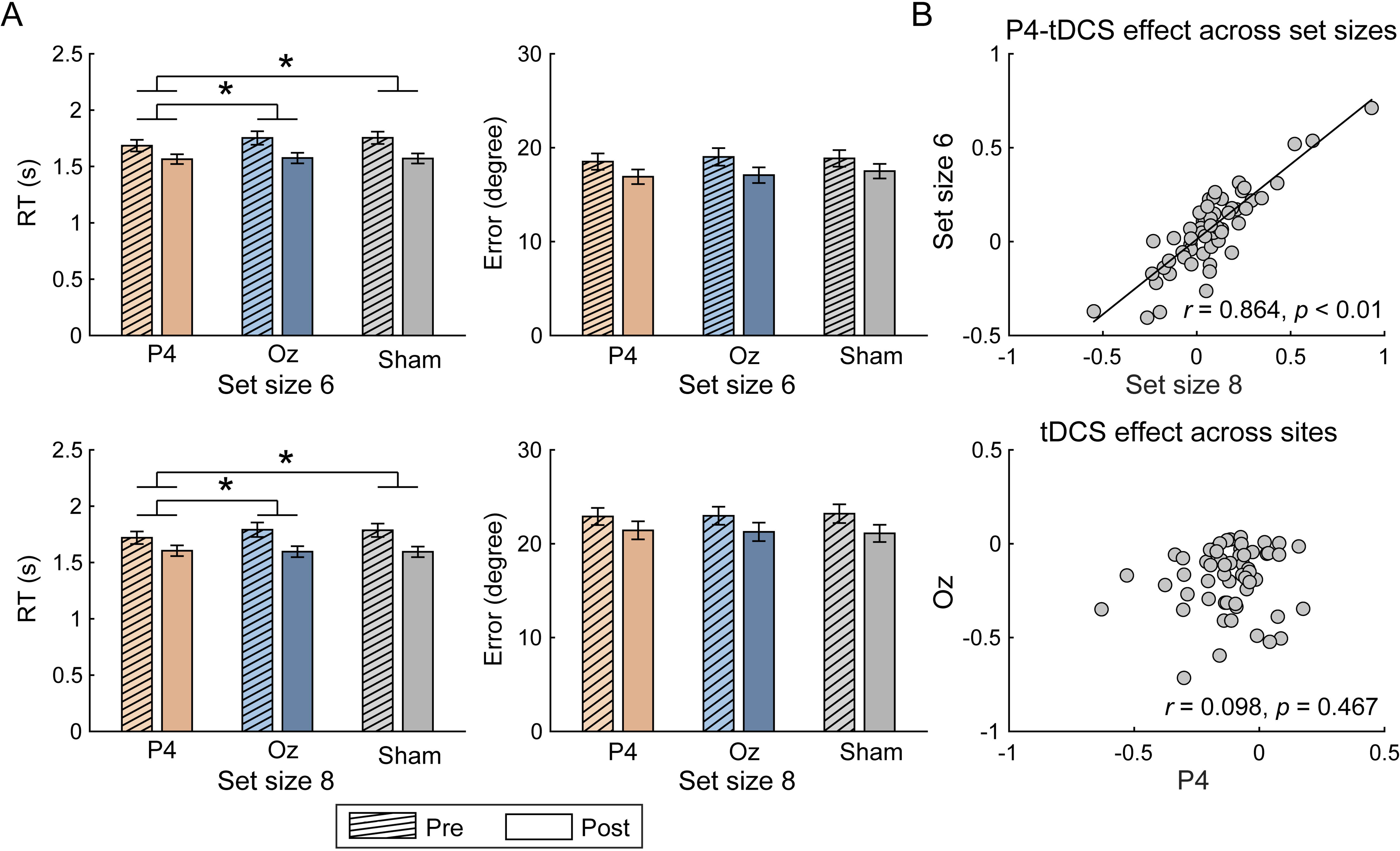
(A) Changes in RT and recall error across stimulation conditions and memory loads. The error bars represent standard error (SEM). Statistical test markers indicate significant stimulated region × test time interaction. * *p* < 0.05. (B) Correlation of tDCS effect over PPC across set sizes (top) and tDCS effect over regions (bottom).

More importantly, we also found significant interactions between two stimulation conditions (i.e., PPC and OCC) and test time for both memory loads (*Fs* < 5.969, *ps* < 0.038, *η_p_*^2^ s < 0.096, BF_10_s < 4.908). Similarly, further analyses found that the RT reduction was significantly smaller after PPC stimulation (set size 6: *t*_(56)_ = 6.071, *p* < 0.001, Cohen’s *d* = 0.804, BF_10_ > 1000; set size 8: *t*_(56)_ = 5.406, *p* < 0.001, Cohen’s *d* = 0.716, BF_10_ > 1000) than that after OCC stimulation (set size 6: *t*_(56)_ = 8.179, *p* < 0.001, Cohen’s *d* = 1.083, BF_10_ > 1000; set size 8: *t*_(56)_ = 7.810, *p* < 0.001, Cohen’s *d* = 1.034, BF_10_ > 1000). However, our results revealed that effect sizes of tDCS over PPC and OCC were not correlated across subjects (*r*_(55)_ = 0.098, *p* = 0.467, BF_10_ = 0.214, Figure 2B, bottom). Together, these results indicated that PPC stimulation selectively prolonged RT.

Meanwhile, we did not find any significant interaction between test time and stimulation condition for tDCS effects on recall errors (*F*s < 2.202, *p*s > 0.143, *η_p_*^2^ s < 0.038, BF_10_s < 0.218; Figure 2A, right), suggesting no overall VWM performance changes after PPC or OCC stimulation.

#### PPC stimulation increased non-target responses

For tDCS effects over PPC on *p*NT, we found significant interaction among stimulation condition, stimulation time, and memory load (*F*_(1,56)_ = 10.226, *p* = 0.002, *η_p_*^2^ = 0.154, BF_10_ = 4.925). Follow-up two-way ANOVA showed that the interaction between the stimulated region and test time was only significant in set size 8 condition (*F*_(1,56)_ = 7.593, *p* = 0.008, *η_p_*^2^ = 0.119, BF_10_ = 3.159; in set size 6: *F*_(1,56)_ = 2.157, *p* = 0.147, *η_p_*^2^ = 0.037, BF_10_ = 0.473; Figure 3A). For set size 8, non-target responses were comparable before and after PPC stimulation (*t*_(56)_ = 0.254, *p* = 0.801, Cohen’s *d* = 0.034, BF_10_ = 0.149) but became significantly lower after sham stimulation (*t*_(56)_ = 3.426, *p* = 0.001, Cohen’s *d* = 0.454, BF_10_ = 24.009). Unlike the tDCS effects on RTs, the effect sizes of tDCS over PPC across memory loads were not correlated (*r*_(55)_ = 0.062, *p* = 0.646, BF_10_ = 0.183; Figure 3B, top).

**Figure 3.**
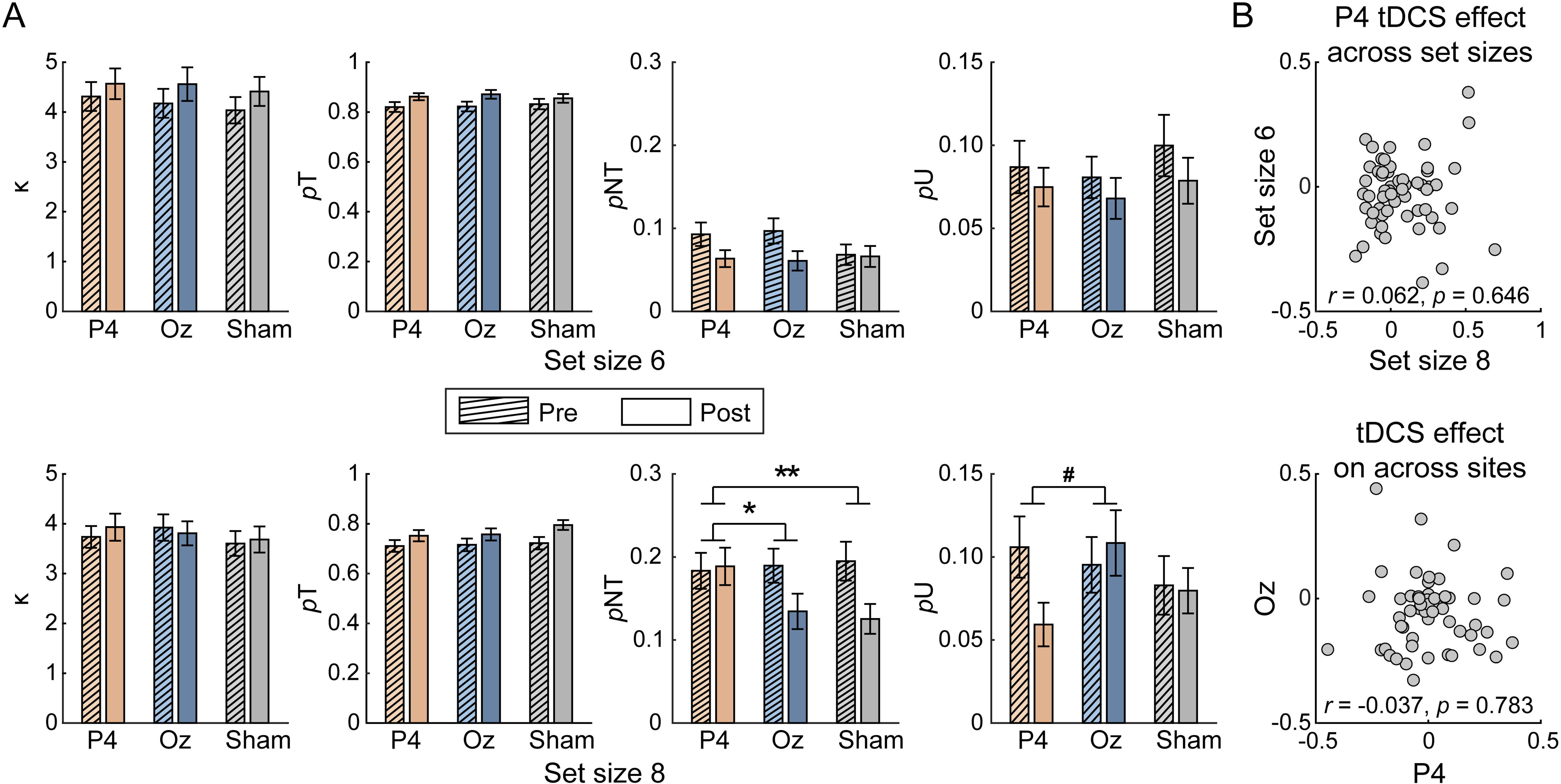
(A) Changes in response precisions (κ), probability of target responses (*p*T), probability of non-target responses (*p*NT), and probability of guessing (*p*U) across different stimulation conditions and memory loads. The error bars represent standard error (SEM). Statistical test markers indicate significant stimulated region × test time interaction. # 0.05 < *p* < 0.1, \**p* < 0.05, \*\**p* < 0.01. (B) Correlation of tDCS effect over PPC across set sizes (top) and tDCS effect over regions (bottom).

To further examine the specificity of the tDCS effects of PPC on *p*NT, we compared *p*NT changes before and after stimulation over PPC and OCC in set size 8. The interaction between the stimulated region and test time was significant (*F*_(1,56)_ = 4.455, *p* = 0.039, BF_10_ = 2.319). Specifically, comparable *p*NTs were observed after PPC stimulation (*t*_(56)_ = 0.254, *p* = 0.801, Cohen’s *d* = 0.034, BF_10_ = 0.149) whereas decreased *p*NTs were found after OCC stimulation (*t*_(56)_ = 2.944, *p* = 0.005, Cohen’s *d* = 0.390, BF_10_ = 6.563). Similarly, there was no correlation between the tDCS effects over PPC and OCC on *p*NT (*r*_(55)_ = -0.037, *p* = 0.783, BF_10_ = 0.172; Figure 3B, bottom). Besides, additional correlation analysis revealed that tDCS effects on RT and *p*NT were also independent (*r*_(55)_ = 0.073, *p* = 0.589, BF_10_ =0.191). Together, these results suggested that, compared with sham stimulation, PPC stimulation specifically increased non-target responses in the high memory load.

Besides, for tDCS effects over PPC on precisions, target probability, and guessing probability, we only found a marginal three-way interaction effect among stimulated region, test time and memory load in *p*U (*F*_(1,56)_ = 3.244, *p* = 0.077, *η_p_*^2^ = 0.055, BF_10_ = 1.542; others: *F*s < 2.732, *p*s > 0.104, *η_p_*^2^ s < 0.047, BF_10_s < 0.159). Further analysis indicated that, in set size 8, PPC stimulation decreased the random guesses (*t*_(56)_ = 2.535, *p* = 0.014, Cohen’s *d* = 0.336, BF_10_ = 2.670) while no such difference in sham stimulation (*t*_(56)_ = 3.426, *p* = 0.001, Cohen’s *d* = 0.454, BF_10_ = 24.009). Similarly, neither three-way interactions nor two-way interactions were observed for OCC stimulation (*F*s < 2.862, *p*s > 0.434, BF_10_s < 0.504), showing no tDCS effect over OCC on these parameters.

#### tDCS effects over PPC were not modulated by capacity or strategy

Correlation results showed that individual capacity or the recall strategy index were not correlated with the PPC tDCS effects of prolonged RT or increased *p*NT (*r*s < -0.213, *p*s > 0.112, BF_10_s < 0.573; Figure 4).

**Figure 4.**
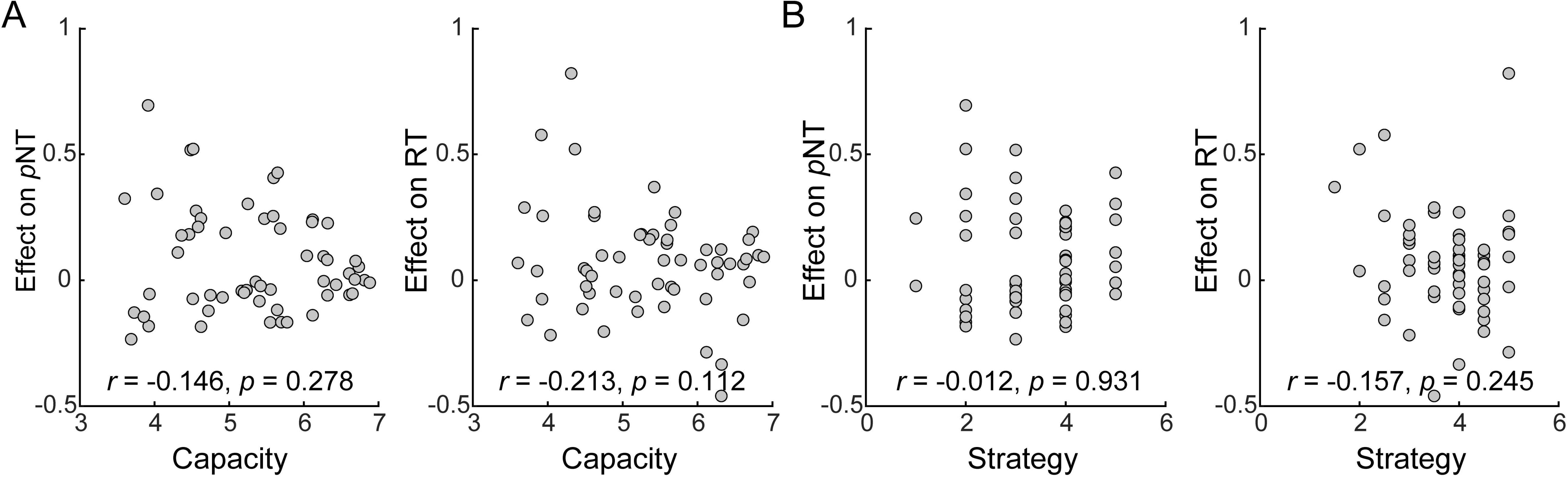
(A) Correlation between memory capacity and tDCS effects. (B) Correlation between memory strategy and tDCS effects. Note that effects on *p*NT were calculated under set size 8, effect on RT was averaged across two memory loads.

#### Summary of Experiment 1

Experiment 1 revealed that PPC stimulation specifically prolonged RT and increased *p*NT in the delayed estimation task, supporting the view that the parietal lobe played a critical role in content-context binding during VWM (Cai et al., 2020; Gosseries et al., 2018). In contrast, although studies reported OCC played an important role in information representation during VWM (Bettencourt & Xu, 2016), stimulation over the occipital lobe did not change VWM performance. The results suggested the higher brain areas may play a more causal role in VWM.

Based on the latest metacognitive model, van den Berg et al. (2017) proposed two types of non-target responses: high-confidence mis-binding and low-confidence informed guessing. In Experiment 2, we further examined whether PPC stimulation equally affected these two types of non-target responses.

## Experiment 2

### Participants

The sample size was determined using G*Power 3.1 (Faul et al., 2007) based on the effect size of increased *p*NT after PPC stimulation in Experiment 1 (*η_p_*^2^ = 0.119). A minimum sample of 28 participants was required to achieve a power of 90%, with a significance level of 0.05. We recruited 33 healthy university students (14 females; *M* = 23.10 years, *SD* = 2.70), and four participants were excluded due to poor task performance, resulting in a final sample of 29 in this experiment. Recruitment and payment criteria were consistent with Experiment 1.

### Experimental procedure and tDCS setup

The task procedure was similar to Experiment 1, except that participants were required to make metacognitive evaluations of their recall during the task. More specifically, in each trial, after 1 s delay, a white circle cueing the location of the probed orientation appeared for 200 ms and participants were asked to report whether they remembered or forgot the probed orientation by pressing the left or right button of the mouse. The left-right buttons were counterbalanced across participants. Subsequently, participants were instructed to recall the targeted orientation at the probed position within 4 s by moving the mouse and making a confirming click. Finally, participants rated their confidence in a 7-point scale (1 = “lowest confidence”; 7 = “highest confidence”), which was introduced as a validation of the remember-forget binary forced choice (Figure 5A).

**Figure 5.**
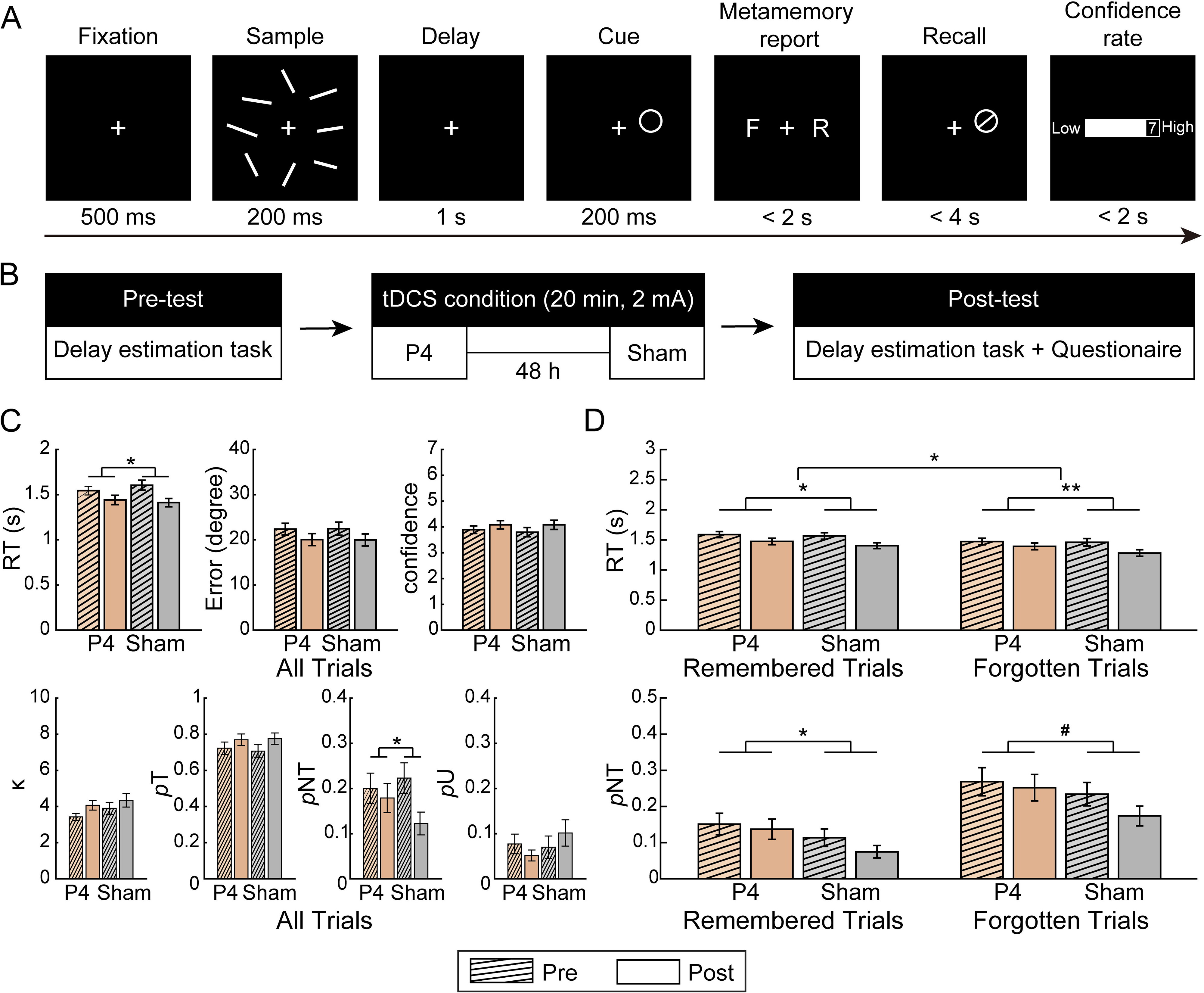
(A) Schematic diagrams of the delay estimation task. (B) tDCS procedures. (C) Changes in RT, error, confidence rating, and fitting parameters across stimulation conditions. The error bars represent standard error (SEM). Statistical test markers indicate significant interactions. # 0.05 < *p* < 0.1, * *p* < 0.05, ** *p* < 0.01. (D) Changes in RT and *p*NT for remembered and forgotten trials across stimulation conditions.

Participants completed six blocks of the delay estimation task before and after stimulation, each block included 60 trials, and each task lasted for approximately 20 minutes. In this experiment, we only focused on the high set size condition (i.e., set size 8) and included two tDCS sessions (i.e., PPC and sham). All stimulation setups were similar to Experiment 1 (Figure 5B).

### Data analysis

#### Replicate the overall tDCS effects on RT and non-target response

Across all trials, we first examined tDCS effects over PPC on prolonged RT and increased *p*NT observed in Experiment 1 by conducting a two-way repeated measures ANOVA for stimulation condition (PPC vs. sham) and test time (pre-test vs. post-test). For significant two-way interaction effects, we further conducted simple effect tests. Similar to Experiment 1, for significant tDCS effects, we explored the potential influences of capacity and strategy through correlation analysis. Here, we used an advanced model fitting method to obtain all three parameters (*p*NT, *p*T, *p*U) for each trial (Schneegans & Bays, 2016).

#### Compare tDCS effects between remembered and forgotten trials

Trials were divided into remembered trials (Trial number: *Mean* = 220.974, *SD* = 12.872) and forgotten trials (*Mean* = 131.371, *SD* = 10.570) based on the binary forced choice. Paired *t*-tests revealed that errors for remembered trials were significantly lower than those for forgotten trials (*Mean (SD) =* 16.280 (5.743) vs. 29.208 (7.630), *t*_(28)_ = -16.174, *p* < 0.001, Cohen’s *d* = -3.003, BF_10_ > 1000), and confidence ratings were also significantly higher for remembered trials than for forgotten trials (*Mean (SD)* = 4.922 (0.960) vs 2.302(0.629), *t*_(28)_ = 15.638, *p* < 0.001, Cohen’s *d* = 2.904, BF_10_ >1000), suggesting that our metacognition categorization was validated and forced choice self-reports reflected objective memory. Then, behavioral parameters for each trial type were averaged for follow-up tDCS effect comparisons.

For the tDCS effects on RT and *p*NT, we examined whether these effects differed across trial types through a three-way repeated measures ANOVA of trial type (remembered vs. forgotten), stimulation condition (PPC vs. sham), and test time (pre-test vs. post-test). We further conducted two-way repeated measures ANOVAs and simple effect tests for stimulation condition and test time in each trial type if the three-way interaction effects were significant. Other parameters were also tested in a similar way.

### Results

#### PPC stimulation prolonged recall RT and increased non-target responses

Consistent with Experiment 1, we observed a significant interaction between stimulation condition and test time on RT (*F*_(1,28)_ = 5.894, *p* = 0.028, *η_p_*^2^ = 0.174, BF_10_ = 5.178). Post-hoc tests revealed that the RT decrease after PPC stimulation (*t*_(28)_ = 4.563; *p*s < 0.001, Cohen’s *d* = 0.847, BF_10_ = 282.944) was significantly smaller than that after sham stimulation (*t*_(28)_ = 7.356; Cohen’s *d* = 1.366, *p*s < 0.001, BF_10_ > 1000). For *p*NT, the interaction between stimulation condition and test time was also significant (*F*_(1,28)_ = 4.498, *p* = 0.045, *η_p_*^2^ = 0.138, BF_10_ = 2.691).

Post-hoc analyses revealed that there was no difference after PPC stimulation (*t*_(28)_ = 0.836, *p* = 0.410, Cohen’s *d* = 0.155, BF_10_ = 0.272) but a significant decrease in *p*NT after sham stimulation (*t*_(28)_ = 3.816, *p* < 0.001, Cohen’s *d* = 0.709, BF_10_ = 46.613). Meanwhile, tDCS effects over PPC on RT and *p*NT were independent across participants (*r*_(27)_ = -0.256, *p* = 0.180, BF_10_ = 0.544) and neither tDCS effect was correlated with individual differences in capacity or strategy (*r*s < - 0.297, *p*s > 0.118, BF_10_s < 0.739). Besides, despite a similar numerical trend of two-way interaction between stimulated region and test time was observed for *p*U (*F*_(1,28)_ = 2.780, *p* = 0.107, *η_p_*^2^ = 0.090, BF_10_ = 1.497), no other effects on recall error, confidence ratings, and other parameters were significant (*F*s < 0.978, *p*s > 0.331, *η_p_*^2^s < 0.034, BF_10_ s < 0.403).

#### tDCS effects over PPC on RT were greater in forgotten trials than in remembered trials

We further examined the tDCS effects over PPC between forgotten and remembered trials. For RT, the interaction among trial type, stimulation condition, and test time was significant (*F*_(1,28)_ = 6.385, *p* = 0.017, *η_p_*^2^ = 0.186, BF_10_ = 2.372; Figure 5D, top). Further two-way repeated-measures ANOVAs for each trial type indicated that the effect size in forgotten trials (*F*_(1,28)_ = 11.506, *p* = 0.002, *η_p_*^2^ = 0.291, BF_10_ = 65.871) was larger than in remembered trials (*F*_(1,28)_ = 4.293, *p* = 0.048, *η_p_*^2^ = 0.133, BF_10_ = 2.244). For *p*NT, however, no significant interaction effect among trial type, stimulation condition, and test time was observed (*F*s < 2.707, *p*s > 0.111, *η_p_*^2^s < 0.088, BF_10_s < 0.225; Figure 5D, bottom), indicating comparable tDCS effects in two types of non-target trials.

#### Summary of Experiment 2

In Experiment 2, we replicated the main findings of Experiment 1 that PPC stimulation increased RT and *p*NT during VWM. More importantly, compared with remembered trials, tDCS effects on RT were greater in forgotten trials while were comparable on *p*NT. In sum, these results suggested that PPC was causally involved in two types of non-target responses, while may through different mechanisms.

Inspired by a recent computational modeling study indicating that VWM recall and recognition involved different cognitive processes (Kahana, 2020). Delayed estimation and change detection are two typical VWM tasks for memory recall and recognition. Therefore, we further examined whether the tDCS over PPC mainly affected VWM retrieval processes and caused behavioral changes, by using a change detection task instead of the delayed estimation task in Experiment 3.

## Experiment 3

### Participants

Thirty-four healthy university students (18 females; *M* = 22.00 years, *SD* = 2.06) were recruited. Six participants were excluded due to poor performance, leaving 28 participants for the following analysis. The sample size choice, participant recruitment, and payment criteria were consistent with Experiment 2.

### Materials and methods

The change detection task for orientations was adapted from a previous study (Gong & Li, 2014; Figure 6A). The encoding and maintenance periods were the same with the delayed estimation task. During the probe, a probed orientation appeared on the screen and participants were required to make a judgment about whether the probed orientation changed compared with the orientation presented in the same location during the sample period, by pressing either the “F” or “J” key. The response keys were counterbalanced across participants and the maximum response window was 2 s. The probabilities of orientation change and no change were equal. In the change trials, half of the probed orientation was different from all the orientations during the sample display (i.e., "change trials") and the other half of the probed orientation was the same as the orientation located next to the probed location (i.e., "lure trials"). The change detection task consisted of 240 trials for each memory load (i.e., set size 6 and set size 8). Before and after stimulation, participants completed four blocks of the task, and each block consisted of 60 trials with two memory loads randomly mixed. The tDCS setup was consistent with Experiment 2 (Figure 6B).

**Figure 6.**
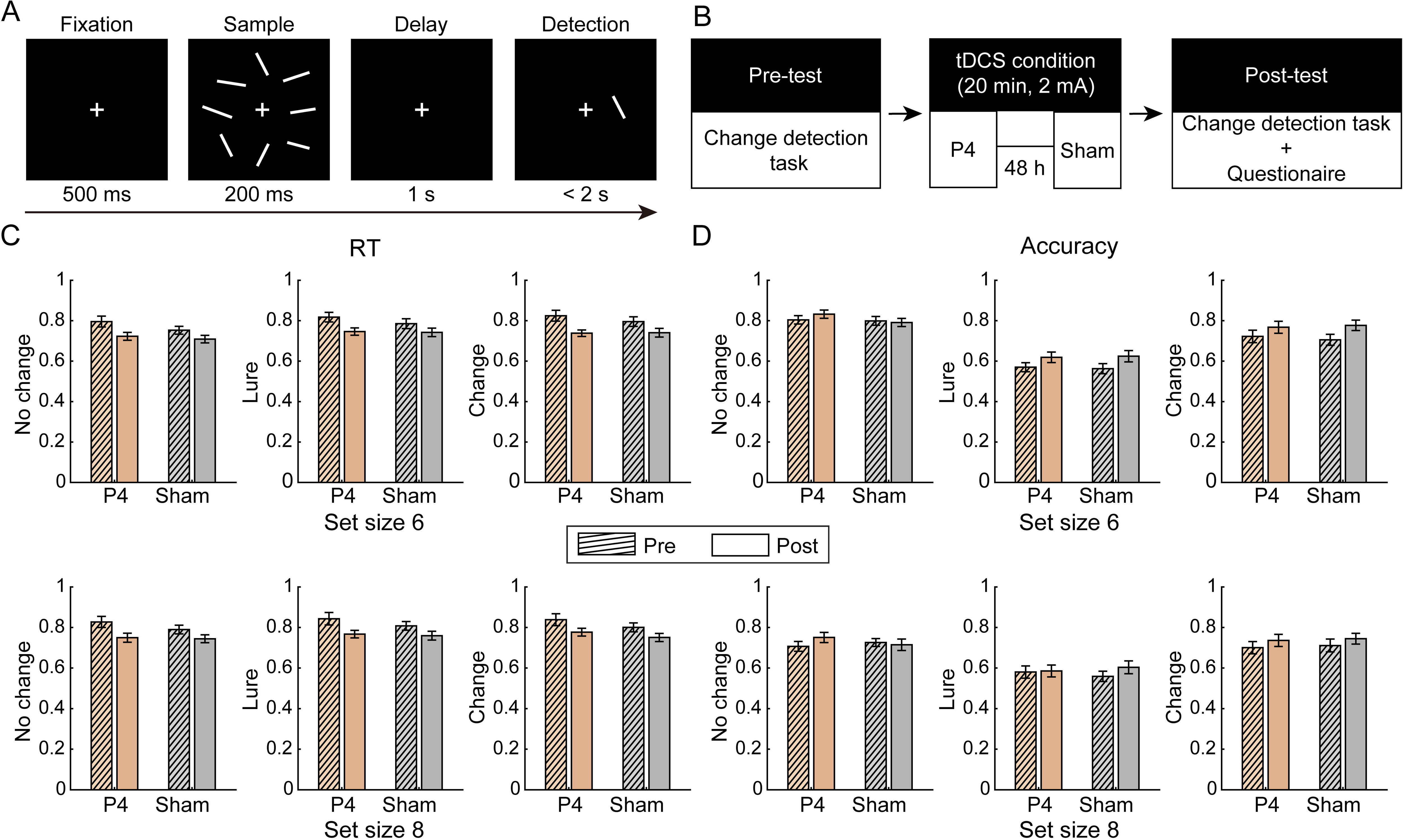
(A) Schematic diagrams of the change detection task. (B) tDCS procedures. Changes in RT (C) and accuracy (D) across trial types and stimulation conditions. The error bars represent standard error (SEM).

### Data analysis

#### Estimation of behavioral performance and tDCS effects

First, we calculated overall RT and accuracy under each memory load based on all trials. RT was defined as the duration between the onset of the probe and button-press response, and accuracy referred to the proportion of correctly response trials out of all trials. Then, we also obtain RT and accuracy in no change trials, lure trials, and change trials, respectively. In lure trials, higher accuracy indicated better feature binding ability and lower non-target responses (e.g., lower *p*NT in Experiment 1 and 2). To examine the tDCS effects on each behavioral parameter, we performed a series of 2 (stimulated region: P4 vs. sham) × 2 (test time: pre vs. post) × 2 (memory load: set size 6 vs. set size 8) ANOVAs. If their interactions were significant, we further analyzed tDCS effects on each memory load. Based on the findings of Experiment 1 and 2, the effects of tDCS over PPC on overall RT and accuracy in lure trials were of most interest.

### Results

#### No tDCS effect over PPC on response time or mis-binding processes in recognition

No interaction effect between stimulation and test time was significant for RT (*F*s < 1.251, *p*s > 0.273, η*p*s^2^ < 0.044, BF_10_s < 0.390) or for accuracy in lure trials (*F*s < 0.832, *p*s > 0.370, BF_10_s < 0.391), indicating that PPC stimulation did not affect RT or binding processes in VWM recognition (Figure 6C–D). Similarly, there was no significant interaction effect on other behavioral parameters (*F*s < 1.235, *p*s > 0.246, η*p*s^2^ < 0.042, BF_10_s < 0.586; except for a trend effect on accuracy in high memory load in no change trials, *F*_(1,27)_ = 3.441, *p* = 0.075, BF_10_ = 0.945).

## Discussion

The current study established the causal relationship between posterior parietal activity and feature binding during VWM retrieval through three tDCS experiments. First, we found that anodal tDCS over PPC selectively increased recall RT as well as non-target responses in the delayed estimation task. Meanwhile, combined with metacognitive evaluations, we clarified these tDCS effects could be observed in both types of non-target responses (i.e., mis-binding and informed guessing). Besides, we further identified that the effects of tDCS over PPC were specific during memory retrieval by demonstrating that such effects were not observed during memory recognition. Together, our findings deepen our understanding of the involvement of PPC in the feature binding during VWM retrieval.

First of all, in two independent samples, we replicated that anodal tDCS over PPC increased non-target responses in the delayed estimation task. These results supported recent fMRI studies that emphasized the close relationship between posterior parietal activity and the feature binding processes during VWM (Cai et al., 2020; Gosseries et al., 2018). Meanwhile, we noticed the increased non-target responses along with a trend of decreased random guesses. That is, tDCS over PPC biased the recall process. Two possibilities could explain these findings. One possibility is that enhanced posterior parietal activity may facilitate the reinstatement of content information from multiple items during retrieval, leading to an increased probability of non-target responses. Supporting this view, Baddeley (2000) proposed the posterior parietal lobe as a core area for the episodic buffer during WM, and Xie et al. (2017) further interpreted PPC as a hub of multisensory information integration, which claimed PPC’s critical role in representing and combining different features. Moreover, our results emphasized that PPC was causally involved during memory retrieval instead of recognition. Comparing the delayed estimation and change detection tasks, previous studies have shown similar neural activity during WM maintenance. For instance, EEG studies have found similar contralateral delay activity (Adam et al., 2018; Cai et al., 2022; Vogel & Machizawa, 2004) and fMRI studies showed similar frontoparietal activity during these two tasks (Cai et al., 2018; Kim, 2019). In contrast, during the response, a recent study suggested that the delayed estimation task is mainly based on detailed retrieval while the change detection task is based on familiarity judgment (Kahana, 2020). Although few studies directly compared the PPC activity between recall and recognition during WM, considerable evidence has supported the critical role of PPC in episodic memory retrieval but not in recognition. For example, studies have revealed that both activation strength (Cabeza et al., 2008; Sestieri et al., 2017; Wagner et al., 2005) and the neural representations in PPC (Xiao et al., 2017) increased during memory retrieval but not during recognition (Dobbins et al., 2003). Currently, our findings confirmed the causal relationship between the PPC and retrieval during VWM, which was similar to those in episodic memory.

Another possibility proposed that enhanced posterior parietal activity may increase cognitive resources and lead to the adoption of a more proactive retrieval strategy, increasing non-target responses. Our results revealed that tDCS over PPC comparably increased both types of non-target responses (i.e., mis-binding and informed guessing). Researchers have proposed that mis-binding responses were generated when participants mis-organized information across different items while informed guessing reflected participants actively make choices from all the memorized information (van den Berg et al., 2017). Consistently, recent studies revealed that mis-binding mainly resulted from less efficient information processing during early encoding or storage (Emrich & Ferber, 2012; Pertzov et al., 2017; Zokaei et al., 2014) whereas informed guessing more likely reflected the different neural activity during late storage or memory retrieval (Huang, 2020; Pratte, 2019). Consequently, both non-target responses required more cognitive effort compared with random guesses. From the source-consumption perspective, the latest study revealed that parietal tDCS could reduce the cumulative fatigue effect during tasks (Hemmerich et al., 2023). Therefore, we could not exclude the possibility that anodal PPC stimulation enabled individuals to generate both more proactive non-target responses during VWM retrieval.

Besides, we also found that PPC stimulation prolonged recall RT, which was in line with both explanations about increased non-target responses above. However, some evidence in the current study suggested that the tDCS over PPC affected recall RT and feature binding through different pathways. For example, we found that tDCS over PPC only increased non-target responses under high memory load, whereas it changed RTs across memory loads and the effects were highly correlated across loads at the individual level. Meanwhile, we observed greater tDCS effects on RTs in informed guessing trials than in mis-binding trials while comparable effects on non-target responses between these two types of trials. Furthermore, tDCS effects on RTs and non-target responses were always independent at the individual level. Supporting these findings, the latest meta-analysis demonstrated that different PPC stimulation patterns enhanced WM accuracy and RT separately (i.e., different frequencies; Wischnewski et al., 2024). However, future studies are needed to further clarify how PPC is differently involved in the recall speed and accuracy of memory retrieval.

Different from the PPC stimulation effects, it is noteworthy that we did not observe any tDCS effects over the occipital cortex on VWM. A set of recent studies found there was a location-specific neural representation in the occipital cortex which requires accurate item-context binding (Fulvio et al., 2023; Teng & Postle, 2024), and some other studies also reported these neural representations predicted non-target responses at the individual level (Cai et al., 2020; van Lamsweerde & Johnson, 2015). We suggested two possible explanations for this inconsistency. First, unlike the sustained activation of PPC during maintenance, the neural representations of the item or its context information did not depend on sustained activation in the occipital cortex (Harrison & Tong, 2009; Riggall & Postle, 2012). Since tDCS is expected to increase the neural activity of specific areas instead of promoting the neural representation directly, tDCS over the occipital cortex could not significantly affect behavior. Second, previous studies have suggested that occipital neural activity is regulated by feedback signals from the frontal and posterior parietal regions (Halgren et al., 2002). Therefore, even if the occipital neural representation is affected by tDCS, the occipital cortex could maintain the memory information efficiently by receiving feedback signals from other brain areas. Together, these results indicated that the occipital activity has no causal effect on VWM performance or feature binding process.

Finally, some limitations need to be paid attention in the current study. First, we did not find that tDCS over PPC improved VWM capacity, which supported some recent null findings (Dumont et al., 2021; Heinen et al., 2016; Jiang et al., 2024; Robison et al., 2017) instead of earlier studies (Li et al., 2017; Wang et al., 2019). Meanwhile, we also did not replicate the correlation between tDCS effects and individual capacity (Hsu et al., 2014; Tseng et al., 2012) or the adoption of remember-subset strategy (Wang et al., 2020). Our findings may not end this controversy, but these findings strongly reminded us that future studies should systematically explore potential factors influencing tDCS effects, combining the high-definition stimulation settings and neuroimaging methods. Second, we found that single-session tDCS only changed the response bias for non-targets but failed to change overall working memory performance. Future studies using transcranial alternative current stimulation (tACS) or randomized noise stimulation (tRNS) could be tested to further improve feature binding efficiency and the overall VWM performance.

In conclusion, the present study demonstrates that enhanced posterior parietal activity prolongs RT in VWM retrieval and increases the probability of binding errors, and these effects are observed in two types of binding errors (i.e., mis-binding and informed guessing). Our findings provide direct evidence of the causal relationship between the posterior parietal cortex and feature binding, deepening our understanding of the neural basis of feature binding in VWM.

## References

Adam, K. C. S., Robison, M. K., & Vogel, E. K. (2018). Contralateral delay activity tracks fluctuations in working memory performance. Journal of Cognitive Neuroscience, 30(9), 1229–1240.

Ashbridge, E., Cowey, A., & Wade, D. (1999). Does parietal cortex contribute to feature binding? Neuropsychologia, 37(9), 999–1004.

Baddeley, A. (2000). The episodic buffer: A new component of working memory? Trends in Cognitive Sciences, 4(11), 417–423.

Bays, P. M., Catalao, R. F. G., & Husain, M. (2009). The precision of visual working memory is set by allocation of a shared resource. Journal of Vision, 9(10), 1–11.

Bettencourt, K. C., & Xu, Y. (2016). Decoding the content of visual short-term memory under distraction in occipital and parietal areas. Nature Neuroscience, 19(1), 150–157.

Braet, W., & Humphreys, G. W. (2009). The role of reentrant processes in feature binding: Evidence from neuropsychology and TMS on late onset illusory conjunctions. Visual Cognition, 17(1-2), 25–47.

Brainard, D. H., & (1997). The Psychophysics Toolbox. Spatial Vision, 10(4), 433–436.

Cabeza, R., Ciaramelli, E., Olson, I. R., & Moscovitch, M. (2008). The parietal cortex and episodic memory: An attentional account. Nature Reviews Neuroscience, 9(8), 613–625.

Cai, Y., Fulvio, J. M., Samaha, J., & Postle, B. R. (2022). Context binding in visual working memory is reflected in bilateral event-related potentials, but not in contralateral delay activity. eNeuro, 9(6), ENEURO.0207-0222.2022.

Cai, Y., Fulvio, J. M., Yu, Q., Sheldon, A. D., & Postle, B. R. (2020). The role of location-context binding in nonspatial visual working memory. eNeuro, 7(6), ENEURO.0430-0420.2020.

Cai, Y., Urgolites, Z., Wood, J., Chen, C., Li, S., Chen, A., & Xue, G. (2018). Distinct neural substrates for visual short-term memory of actions. Human Brain Mapping, 39(10), 4119–4133.

Dedoncker, J., Brunoni, A. R., Baeken, C., Vanderhasselt, M.-A., & Brunoni, A. R. (2016). A systematic review and meta-analysis of the effects of transcranial direct current stimulation (tDCS) over the dorsolateral prefrontal cortex in healthy and neuropsychiatric samples: Influence of stimulation parameters. Brain Stimulation, 9(4), 501–517.

Dobbins, I. G., Rice, H. J., Wagner, A. D., & Schacter, D. L. (2003). Memory orientation and success: separable neurocognitive components underlying episodic recognition. Neuropsychologia, 41(3), 318–333.

Dumont, R., Majerus, S., & Hansenne, M. (2021). Transcranial direct current stimulation (tDCS) over the intraparietal sulcus does not influence working memory performance. Psychologica Belgica, 61(1), 200–211.

Emrich, S. M., & Ferber, S. (2012). Competition increases binding errors in visual working memory. Journal of Vision, 12(4), 12.

Faul, F., Erdfelder, E., Lang, A.-G., & Buchner, A. (2007). G*Power 3: A flexible statistical power analysis program for the social, behavioral, and biomedical sciences. Behavior Research Methods, 39(2), 175–191.

Fulvio, J. M., Yu, Q., & Postle, B. R. (2023). Strategic control of location and ordinal context in visual working memory. Cerebral Cortex, 33(13), 8821–8834.

Gong, M., & Li, S. (2014). Learned reward association improves visual working memory. Journal of Experimental Psychology, 40(2), 841–856.

Gosseries, O., Yu, Q., LaRocque, J. J., Starrett, M. J., Rose, N. S., Cowan, N., & Postle, B. R. (2018). Parietal-occipitalinteractions underlying control- and representation-related processes in working memory for nonspatial visual features. The Journal of Neuroscience, 38(18), 4357–4366.

Halgren, E., Boujon, C., Clarke, J., Wang, C., & Chauvel, P. (2002). Rapid distributed fronto-parieto-occipital processing stages during working memory in humans. Cereb Cortex, 12(7), 710–728.

Harrison, S. A., & Tong, F. (2009). Decoding reveals the contents of visual working memory in early visual areas. Nature, 458(7238), 632–635.

Heinen, K., Sagliano, L., Candini, M., Husain, M., Cappelletti, M., & Zokaei, N. (2016). Cathodal transcranial direct current stimulation over posterior parietal cortex enhances distinct aspects of visual working memory. Neuropsychologia, 87, 35–42.

Hemmerich, K., Lupiáñez, J., Luna, F. G., & Martín-Arévalo, E. (2023). The mitigation of the executive vigilance decrement via HD-tDCS over the right posterior parietal cortex and its association with neural oscillations. Cerebral Cortex, 33(11), 6761–6771.

Hill, A. T., Fitzgerald, P. B., & Hoy, K. E. (2016). Effects of anodal transcranial direct current stimulation on working memory: A systematic review and meta-analysis of findings from healthy and neuropsychiatric populations. Brain Stimulation, 9(2), 197–208.

Hsu, T.-Y., Tseng, P., Liang, W.-K., Cheng, S.-K., & Juan, C.-H. (2014). Transcranial direct current stimulation over right posterior parietal cortex changes prestimulus alpha oscillation in visual short-term memory task. NeuroImage, 98, 306–313.

Huang, L. (2020). Distinguishing target biases and strategic guesses in visual working memory. *Attention*, Perception & Psychophysics, 82(3), 1258–1270.

Jiang, S., Jones, M., & von Bastian, C. C. (2023). Mechanisms of cognitive change: Training improves the quality but not the quantity of visual working memory representations. Journal of Cognition, 6(1), 42.

Jiang, S., Jones, M., & von Bastian, C. C. (2024). TDCS over PPC or DLPFC does not improve visual working memory capacity. Communications Psychology, 2(1).

Kahana, M. J. (2020). Computational Models of Memory Search. Annual Review of Psychology, 71, 107–138.

Kim, H. (2019). Neural activity during working memory encoding, maintenance, and retrieval: A network-based model and meta-analysis. Human Brain Mapping, 40(17), 4912–4933.

Kirova, A.-M., Bays, R. B., & Lagalwar, S. (2015). Working memory and executive function decline across normal aging, mild cognitive impairment, and Alzheimer’s disease. BioMed Research International, 2015, 748212.

Lee, C., Jung, Y.-J., Lee, S. J., & Im, C.-H. (2017). COMETS2: An advanced MATLAB toolbox for the numerical analysis of electric fields generated by transcranial direct current stimulation. Journal of Neuroscience Methods, 277, 56–62.

Li, S., Cai, Y., Liu, J., Li, D., Feng, Z., Chen, C., & Xue, G. (2017). Dissociated roles of the parietal and frontal cortices in the scope and control of attention during visual working memory. NeuroImage, 149, 210–219.

Makovski, T., & Lavidor, M. (2014). Stimulating occipital cortex enhances visual working memory consolidation. Behavioural Brain Research, 275, 84–87.

Mallett, R., Lorenc, E. S., & Lewis-Peacock, J. A. (2022). Working memory swap errors have identifiable neural representations. Journal of Cognitive Neuroscience, 34(5), 776–786.

Mayer, J. S., Fukuda, K., Vogel, E. K., & Park, S. (2012). Impaired contingent attentional capture predicts reduced working memory capacity in schizophrenia. PloS One, 7(11), e48586.

Oberauer, K., & Lin, H.-Y. (2017). An interference model for visual and verbal working memory. Psychological Review, 124(1), 21–59.

Pertzov, Y., Manohar, S., & Husain, M. (2017). Rapid forgetting results from competition over time between items in visual working memory. Journal of Experimental Psychology, 43(4), 528–536.

Pratte, M. S. (2019). Swap errors in spatial working memory are guesses. Psychonomic Bulletin & Review, 26(3), 958–966.

Riggall, A. C., & Postle, B. R. (2012). The Relationship between Working Memory Storage and Elevated Activity as Measured with Functional Magnetic Resonance Imaging. The Journal of Neuroscience, 32(38), 12990–12998.

Robison, M. K., McGuirk, W. P., & Unsworth, N. (2017). No evidence for enhancements to visual working memory with transcranial direct current stimulation to prefrontal or posterior parietal cortices. Behavioral Neuroscience, 131(4), 277–288.

Salehinejad, M., Kuo, M., & Nitsche, M. (2019). The impact of chronotypes and time of the day on tDCS-induced motor cortex plasticity and cortical excitability. Brain Stimulation, 12(2), 421.

Salehinejad, M. A., Ghanavati, E., Kuo, M.-F., & Nitsche, M. A. (2023). The role of circadian preferred time of day and sleep pressure in tDCS-induced neuroplasticity and associated cognition. Brain Stimulation, 16(1), 203–204.

Salehinejad, M. A., Wischnewski, M., Ghanavati, E., Mosayebi-Samani, M., Kuo, M. F., & Nitsche, M. A. (2021). Cognitive functions and underlying parameters of human brain physiology are associated with chronotype. Nature Communications, 12(1), 4672.

Schneegans, S., & Bays, P. M. (2016). No fixed item limit in visuospatial working memory. Cortex, 83, 181–193.

Sestieri, C., Shulman, G. L., & Corbetta, M. (2017). The contribution of the human posterior parietal cortex to episodic memory. Nature reviews Neuroscience, 18(3), 183–192.

Teng, C., & Postle, B. R. (2024). Investigating the roles of the visual and parietal cortex in representing content versus context in visual working memory. eNeuro, 11(2), ENEURO.0270-0220.2024.

Todd, J. J., & Marois, R. (2004). Capacity limit of visual short-term memory in human posterior parietal cortex. Nature, 428(6984), 751–754.

Tseng, P., Hsu, T. Y., Chang, C. F., Tzeng, O. J., Hung, D. L., Muggleton, N. G., Walsh, V., Liang, W. K., Cheng, S. K., & Juan, C. H. (2012). Unleashing potential: transcranial direct current stimulation over the right posterior parietal cortex improves change detection in low-performing individuals. The Journal of Neuroscience, 32(31), 10554–10561.

van den Berg, R., Yoo, A. H., & Ma, W. J. (2017). Fechner’s law in metacognition: A quantitative model of visual working memory confidence. Psychological Review, 124(2), 197–214.

van Lamsweerde, A., & Johnson, J. (2015). The role of the occipital cortex in capacity limits and precision of visual working memory. Journal of Vision, 15(12), 661.

Vogel, E. K., & Machizawa, M. G. (2004). Neural activity predicts individual differences in visual working memory capacity. Nature, 428(6984), 748–751.

Wagner, A. D., Shannon, B. J., Kahn, I., & Buckner, R. L. (2005). Parietal lobe contributions to episodic memory retrieval. Trends in Cognitive Sciences, 9(9), 445–453.

Wang, S., Itthipuripat, S., & Ku, Y. (2019). Electrical stimulation over human posterior parietal cortex selectively enhances the capacity of visual short-term memory. The Journal of Neuroscience, 39(3), 528–536.

Wang, S., Itthipuripat, S., & Ku, Y. (2020). Encoding strategy mediates the effect of electrical stimulation over posterior parietal cortex on visual short-term memory. Cortex, 128, 203–217.

Wischnewski, M., Berger, T. A., Opitz, A., & Alekseichuk, I. (2024). Causal functional maps of brain rhythms in working memory. Proc Natl Acad Sci U S A, 121(14), e2318528121.

Xiao, X., Dong, Q., Gao, J., Men, W., Poldrack, R. A., & Xue, G. (2017). Transformed neural pattern reinstatement during episodic memory retrieval. The Journal of Neuroscience, 37(11), 2986–2998.

Xie, Y., Xu, Y., Bian, C., & Li, M. (2017). Semantic congruent audiovisual integration during the encoding stage of working memory: an ERP and sLORETA study. Scientific Reports, 7(1), 5112.

Zhang, W., & Luck, S. J. (2008). Discrete fixed-resolution representations in visual working memory. Nature, 453(7192), 233–235.

Zokaei, N., Heider, M., & Husain, M. (2014). Attention is required for maintenance of feature binding in visual working memory. Quarterly Journal of Experimental Psychology, 67(6), 1191–1213.

